# Identifying important interaction modifications in ecological systems

**DOI:** 10.1101/228874

**Authors:** J. Christopher D. Terry, Michael B. Bonsall, Rebecca J. Morris

## Abstract

Trophic interaction modifications, where a consumer-resource link is affected by additional species, are widespread and significant causes of indirect effects in ecological networks. The sheer number of potential interaction modifications in ecological systems poses a considerable challenge, making prioritisation for empirical study essential. Here, we introduce measures to quantify the topological relationship of individual interaction modifications relative to the underlying network. We use these, together with measures for the strength of trophic interaction modifications to identify modifications that are most likely to exert significant effects on the dynamics of whole systems. Using a set of simulated food webs and randomly distributed interaction modifications, we test whether a subset of interaction modifications important for the local stability and direction of species responses to perturbation of complex networks can be identified. We show that trophic interaction modifications affecting interactions with a high biomass flux, those that connect species otherwise distantly linked, and those where high trophic-level species modify to interactions lower in the web have particular importance for dynamics. In contrast, the centrality of modifications in the network provided little information. This work demonstrates that analyses of interaction modifications can be tractable at the network scale and highlights the importance of understanding the relationship between the distributions of trophic and non-trophic effects.

## Introduction

Ecological communities are held together by networks of interactions between populations of different species. The study of the dynamics of whole ecological communities is dominated by trophic interactions (Dunne and Pascual 2006). However, it is increasingly recognised that non-trophic effects must be considered to improve our understanding of the dynamics of ecosystems (Kéfi *et al.* 2012). Interaction modifications (Wootton 1993), where third-party species influence the strength of interactions, are a significant cause of non-trophic effects and are known to be widespread in ecological communities (Werner and Peacor 2003, Kéfi *et al.* 2015). By grouping the diverse processes that can be represented as modifications of functional responses, the trophic interaction modification (TIM) approach (Golubski and Abrams 2011, Terry *et al.* 2017) provides a route to investigate a significant portion of non-trophic effects.

Studies in small empirical model systems have demonstrated that the impact of TIMs can be considerable, comparable with trophic interactions (Schmitz *et al.* 2004), and capable of driving community dynamics (van Veen *et al.* 2005). At the ecosystem level, interaction modifications have often been identified as the cause of unexpected responses to perturbations (Doak *et al.* 2008, Peckarsky *et al.* 2008, Tack *et al.* 2011). However, despite growing interest (Ohgushi *et al.* 2012, Levine *et al.* 2017), applications to complex natural systems have been slower to develop (Kéfi *et al.* 2012). Since the number of possible interaction modifications rises rapidly with increasing community size, identification and prioritisation of the most important interaction modifications (Aufderheide *et al.* 2013) is necessary to accelerate the improvement of our understanding of ecological networks.

Insights from network-based analyses can bring order to highly complex ecological systems. Unfortunately, interaction modifications do not fit directly within conventional network theory (Newman 2010) where species (nodes) are linked in a pairwise manner by interactions (edges) as interaction modifications are inherently multispecies processes (Terry *et al.* 2017). Approaches to deal with this discontinuity within the framework of existing network theory have generally taken two approaches. Most commonly, studies focus on resultant pairwise effects, drawing conventional ‘non-trophic interaction’ links between the modifier and both interactors (Kéfi *et al.* 2012). These systems can be analysed as multi-layer networks (Pilosof *et al.* 2017) keeping non-trophic and trophic networks distinct, however they do not capture the distinctive dynamics of interaction modifications where the strength of each pairwise effect is dependent on the density of the third species. A second approach uses hyper-graphs can be used where interactions can link any number of species simultaneously (Golubski *et al.* 2016), but this method is non-directional and challenging to quantify and as such is unsuited to dynamical analysis.

In this paper we address the question of where, given limited information, we could most profitably start to incorporate interaction modifications into our understanding of community dynamics. To this end, we develop the framework of trophic interaction modifications by introducing analogues of conventional network metrics that can describe how individual interaction modifications are situated within the wider trophic network. We then use these measures, along with previously described measures for the strength of modifications, to highlight features of interaction modifications that can be used to identify those that are particularly important for whole-community level dynamics, thereby aiding the prioritisation of empirical work on those processes likely to have a particular impact.

## Methods

To act as a testbed for the impact of interaction modifications we generated a set of artificial communities parameterised using well established allometries to represent generalised food webs. We represented the relationships between species in an interaction matrix, **A** (also termed a community or Jacobian matrix (Novak *et al.* 2016)) at an assumed equilibrium. Each element details the change in the growth rate of species *i* in response to a small change in the biomass density of species *j*. Hence 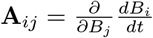, where *B_i_* is the biomass density of species *i*, evaluated at a fixed (equilibrium) point. These interaction matrices were based upon underlying functional response models that were parameterised using empirically derived allometries, to which we added a random set of TIMs as detailed below. All analyses were carried out using R v.3.4.3 (R Core Team 2017) and all code is available in an online repository.

### Model Specification

#### Generating trophic topology and species densities

Trophic network topologies were generated using the niche model algorithm (Williams and Martinez 2000). This simple algorithm has been generally validated as a useful approach to generate networks that have similar structural properties to natural systems (Williams and Martinez 2008). To represent a moderately sized community, we set species richness and trophic connectance at 25 and 0.1 respectively. Cannibalistic interactions were removed and trophic levels were assigned from the network structure following the unweighted pathway method of Levine (1980). Species body-masses, *m_i_*, were assigned based on trophic level following empirical distributions of consumer-resource ratios (Brose *et al.* 2006 a, detailed in SI 1).

The biomass density of each species, *B_i_*, was drawn from a distribution based on the individual body mass as 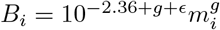 following the approach of Tang *et al.* (2014). The scaling factor between body mass and biomass-density, *g*, can vary considerably between communities (Reuman *et al.* 2009). We therefore replicate our analyses across three values of *g* within the range of empirical estimates (0.25, 0.1 and −0.1). We introduced noise (*ϵ*) to the exponent with mean 0 and standard deviation 0.2 to introduce species-level variation from the precise relationship. To ease comparison between different scaling terms, the mean population density across each community was standardised to 1.

#### Trophic interactions

To model trophic interactions between populations, we used body-mass allometries (Yodzis and Innes 1992, Woodward *et al.* 2005) to parameterise an underlying Type I functional response model, where the rate of loss to a resource population *B_i_* due to a consumer population *B_j_* is *a_ij_B_i_B_j_* and the corresponding rate of gain of the consumer is *e_ij_a_ij_B_j_B_i_*. Biomass-density specific attack rates, *a_ij_*, were defined based on the documented relationship between consumer-resource size ratios and per-capita attack rates (Pawar *et al.* (2012) incorporating a penalty for consumer generality, detailed in SI 1). Conversion efficiencies, *e_ij_*, were drawn from a uniform distribution ranging from 0.1 to 0.2. Off-diagonal elements of the community matrix **A** representing the trophic effect of an increase of the population of a consumer *j* on the growth rate of resource *i* are therefore **A**_*ij*_ = −*a_ij_B_i_* while the corresponding reciprical effect **A**_*ji*_ is given by *e_ij_a_ij_B_j_*. Since we focus our analyses on interspecific interactions we make the implicit assumption that the trophic interactions are balanced by single species processes including intrinsic growth rates, mortality and other extrinsic processes to create an equilibrium with the requisite overall zero net rate of change at the specified biomass densities (see SI 8 for further justification).

#### Self-regulation terms

The diagonal terms of an interaction matrix, **A**_*ii*_, represent the change in the growth rate of each population in response to a small increase in that population. In biological terms they represent density-dependent self-regulation terms, but are determined by a complex mixture of trophic and non-trophic processes (see SI 8). There is no empirical consensus about the strength and distribution of self-regulation in natural systems, but it is likely that self-regulatory negative values are prevalent (Barabás *et al.* 2017).

We therefore directly specify the diagonal terms of the interaction matrix with two separate approaches to confirm that our results are not dependent on particular assumptions. In the first approach (i), each species was assigned a negative **A**_*ii*_ term equal to twice the mean strength of the effects (trophic and non-trophic) exerted upon it. In a second approach (ii), producers were assumed to be more strongly self-regulating than consumers. Each producer was assigned terms **A**_*ii*_ equal to twice the strongest trophic interaction in the network while each consumer was assigned a term 100 times smaller. For both approaches, these values were chosen to calibrate the self-regulation to the trophic interactions while introducing self-regulation at a level where the local stability of populations is likely but not inevitable.

#### Interaction modifications

We model interaction modifications by introducing a relationship between the density of the modifier species, *B_k_* and the post-modification attack rate 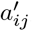:

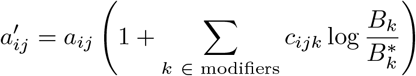

As the density of the modifier species *B_k_* diverges from its equilibrium value 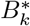, the size and direction of change is specified with a slope parameter, *c_ijk_*. A positive *c_ijk_* causes an increased level of the modifier to strengthen the underlying trophic interaction. A negative value would lead to a weakening of the interaction.

This model formulation, and dependence on the divergence from 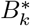 rather than the value of *B_k_*, means that the model assumes that the trophic interaction is already at ‘post-modification’ strength. Hence the allometrically specified interaction strengths are assumed to incorporate the influence of the modifier at the equilibrium density. This allows the effect of interaction modifications to be clearly distinguished from disruption of the trophic network, something that has not been possible in previous dynamic analyses of the effect of interaction modifications on system dynamics (e.g. Arditi *et al.* 2005, Goudard and Loreau 2008, Lin and Sutherland 2013) where the introduction of interaction modification shifts the underlying trophic interaction distribution.

Since the population grwoth rate of both *B_i_* and *B_k_* is now dependent on the density of species *B_k_*, each TIM causes two non-trophic effects, from the modifier species to each of the pair of trophic interactors. These impacts have also previously be described as ‘trait-mediated indirect effects’ (Abrams 2008, Okuyama and Bolker 2012) amongst others. We incorporate these effects into the interaction matrix through the addition of two additional terms, found by taking the derivative of the modified functional response terms with respect to the modifier species:

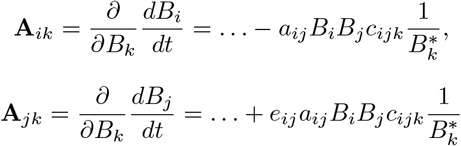

where the ‘…’ indicate trophic terms and other non-trophic effects. Species are assumed to not modify their own trophic interactions (*k ≠ j* or *i*), or alternatively any self-modification is considered subsumed into the self-regulatory terms. Each potential TIM (combination of consumer, resource and modifier species) had an equal and independent probability (0.05) of being present. This resulted in an expectation of 69 TIMs per community of 25 species and trophic connectance 0.1. Slope parameters (*c_ijk_*) for each extant TIM were drawn from a uniform distribution ranging from −0.5 to 0.5. When isolated, the resultant log-normal distribution of non-trophic effect components largely overlapped that of the trophic interactions (SI. 2), in line with available results from meta-analyses (Bolnick and Preisser 2005, Preisser *et al.* 2005).

### Quantifying TIM Topology

TIMs form a distinct class of the connections between species, directly linking species and interactions (Terry *et al.* 2017). Their position within the community can be quantified with reference to the position of the modifier, consumer and resource species within the underlying trophic network. Here, we use seven aspects of topology to describe how individual TIMs are located in the network (Fig. 1). We calculate the trophic level of each species using the unweighted trophic network and the pathway method of Levine (1980). The trophic height of the modifier species and of the interaction (calculated as the mean trophic level of the consumer and the resource species) provide basic information about the position of the modification in food web. These can then be compared to measure what we term the ‘trophic direction’ of the TIM which we define as the trophic height of the interaction minus the trophic level of the modifier. Therefore, positive values imply modifications to interactions higher in the web than the modifier and negative values imply the opposite. The ‘trophic span’ of a modification constitutes the mean number of trophic links from the modifying species to each interactor by the shortest route and measures the extent to which a TIM links otherwise disconnected species.

**Figure 1:**
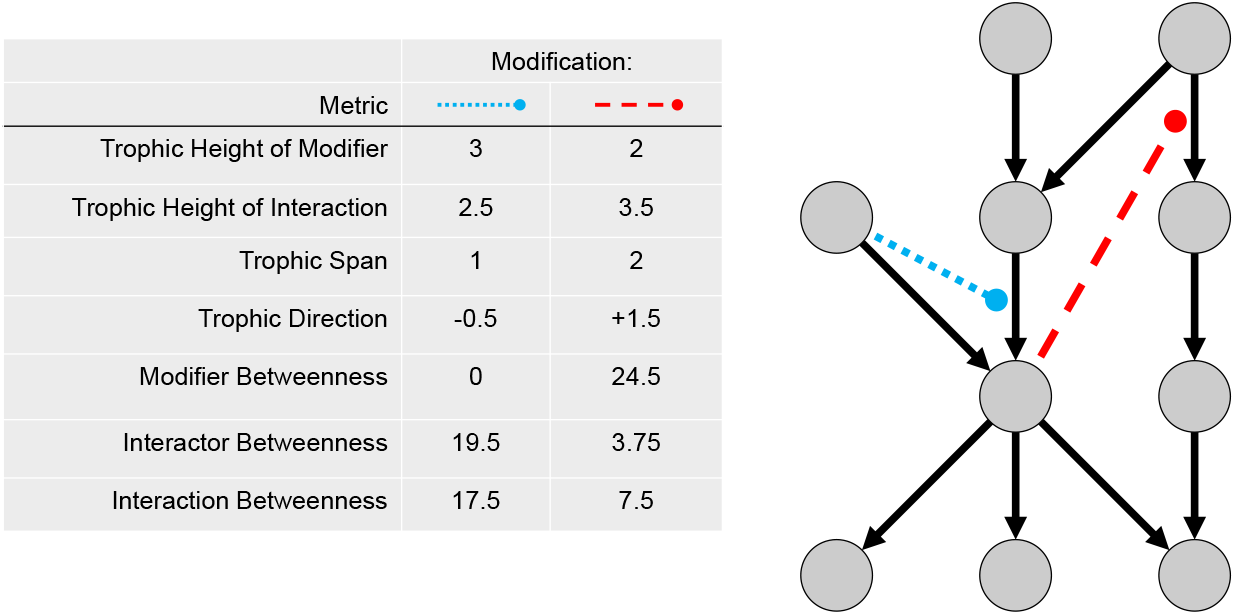
Example values of topological metrics for interaction modifications (dashed lines) within a simple trophic network (solid lines).

The centrality of a TIM can be used to determine the connectivity of the TIM to the wider trophic network - does it involve peripheral species or those that are closely linked to many other species? Centrality of a node or an edge within a network can be quantified by a wide diversity of algorithms (Newman 2010). Here we use ‘betweenness’, a relatively non-technical measure that counts the number of shortest paths between all nodes that pass through a given node or interaction. We only consider unweighted trophic links (excluding TIM connectance) when calculating betweeness as this information is more widely available. We measure the ‘modifier betweeness’, ‘interactor betweenness’ (the mean of the centrality of the two interactors) and ‘interaction betweenness’ (the centrality of the interaction being modified).

### Quantifying the Strength of TIMs

Determining the strength of a TIM is not straightforward (Terry *et al.* 2017). In a similar manner to pairwise interaction strengths (Laska and Wootton 1998, Berlow *et al.* 2004) they can be simultaneously considered with multiple useful approaches. The most basic analyses consider features of the modifier species and the modified interaction. Here, we test the information contained in the density of the modifier species *B_k_*, the size of the underlying trophic interaction as measured by the attack rate (*a_ij_*), and the rate of biomass flux, which in our linear model is *a_ij_B_i_B_j_*.

In our linearised equilibrium systems we directly measure the strength of a modification in four distinct ways. Firstly, we use the ‘slope parameter’ *c_ijk_* as a direct measure of the strength of the relationship between the modifier species and the interaction. Secondly, we determine the strength of the non-trophic effect (‘trait-mediated indirect effect’) as calculated through Eq.XX. Thirdly, we assess the impact on the interaction itself, taking the derivative of the interaction matrix element with respect to the modifier species: 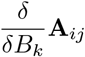. In our model this is determined as 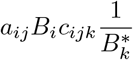. Fourthly, we examine the effect of the TIM on the ‘net-effect’ of the consumer on the resource, taking into account all interaction chains. We calculate this numerically, finding the derivative of **A**^−1^ with respect to a change in the density of the modifier within the TIM. Note that in the latter three cases we solely consider the effect of the consumer on the resource, since in our simple model the reverse is very closely related.

### Independence of Metrics

In general the ‘strength’ properties of TIMs were correlated with each other but topological features considerably less so (Figure 4). Interaction flux was negatively correlated with interaction trophic level despite a slightly positive (0.1) body mass and biomass-density, relationship *g*. In this case the increase in the biomass-density of larger higher-trophic level species was counteracted by reduced attack rates from higher consumer resource body-mass ratios at higher trophic levels.

### Testing Features of Important IMs

We assessed the ability of metrics to identify influential TIMs by testing how well a system containing only a subset of interaction modifications that had been selected using a particular feature was able to capture the dynamics of the complete system (Figure 2). We assess dynamics in two ways discussed below: 1) accuracy in estimating local stability and 2) the capacity to determine the direction of species responses to sustained perturbations, which we will refer to as ‘directional determinancy’.

**Figure 2:**
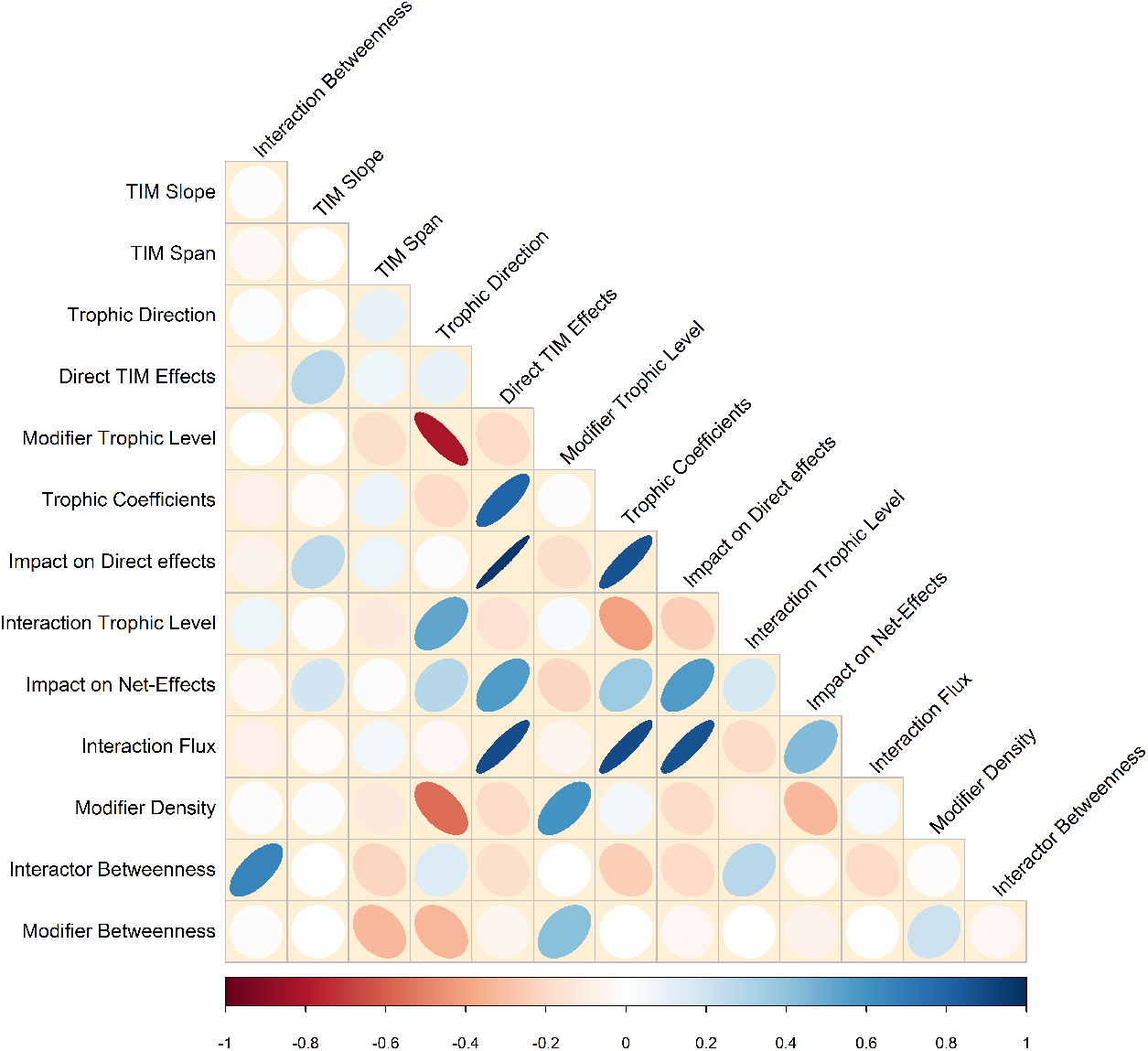
Correlation plot showing relationships between the properties of the TIMs in a model food web community (*g* = 0.1, self-regulation included here by the approach i) described in the main text. Ellipses depict Spearman’s rank correlations, blue shapes indicating positive correlations and red negative. Corresponding plots for the other model structures are shown in SI 6.

#### Stability

Local asymptotic stability is an extensively studied dynamic system property (May 1973, Allesina and Tang 2012). It determines whether a system will (eventually) return to its previous equilibrium after a small perturbation to a population. It can be determined from the sign of the real part of the largest eigenvalue of a system’s community matrix, 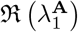. Where this is negative the system can be considered locally stable. With sufficient self-regulation any system can be stabilised. To focus our analysis on the impact of the interaction modifications, rather than the distribution of self-regulation terms (van Altena *et al.* 2016, Barabás *et al.* 2017), for the purposes of the calculation of stability we set each **A**_*ii*_ to 0. In this case, 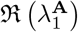 is always positive but can be interpreted as a degree of the self-regulation that would be necessary to stabilise the community. For brevity we will refer to this quantity as ‘stability’.

We generated 2000 communities with interaction modifications for each value of biomass density scaling factor *g*. For each web, the TIM metrics described in the previous section were calculated for each TIM and the ‘top’ 20% of TIMs were identified. Where there was no obvious expectation about which extreme would be more important, for example ‘centrality’, both the highest and lowest values were tested independently. A random subset of 20% of TIMs was also generated for each web to establish a baseline to test metrics against. For each web and each subset of TIMs, a new community matrix **S** was calculated (Figure 2). We then calculated the magnitude of the difference in stability between the full and subset model as:

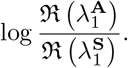

The statistical significance of a metric’s ability to identify a subset of TIMs that estimate stability differently to a random subset of TIMs was tested with a Wilcoxon signed rank test, paired by the underlying community.

#### Direction of response to perturbation

The negative inverse of a community matrix (−**A**^−1^) encapsulates the responses (via all pathways, interaction chains and interaction modifications) of each species to a sustained, small, perturbation to each other species (Bender *et al.* 1984) to generate a ‘net-effects matrix’. This relies on the assumption that the perturbation is sufficiently small that any non-linearities in the response are well represented by the linear approximation of the underlying community matrix.

While estimates of the exact values of responses would be empirically unfeasible, the sign of the net-effects between species, i.e. whether a change in a particular population will lead to an increase or a decrease in another population is a reasonable objective (Yodzis 1988, Novak *et al.* 2011, Aufderheide *et al.* 2013).

We tested the proportion of relationships between species where the sign of the net-effects matrix of the full model −**A**^−1^ matches that of the model containing only a subset of the TIMs −**S**^−1^. This ‘directional determinacy’, also termed ‘predictability’ (Novak *et al.* 2011, Iles and Novak 2016) is a measure of the capacity of a reduced model to represent the dynamics of a full model. It therefore ranges from a minimum of approximately 0.5, where the subset model is no better than chance at correctly identifying species responses, to 1 where the subset model is able to perfectly predict the ‘true’ response. Analytic expressions of the impact on directional determinacy due to the misspecification of Jacobian matrices have recently been derived (Koslicki and Novak 2018). However, since for multiple error terms the expressions become very complex, we directly calculate the effect numerically.

Because local asymptotic stability is a prerequisite for calculating directional determinacy, additional communities were generated and unstable communities excluded to create 2000 locally stable communities for each of the three values of *g* and the two approaches to including self-regulation terms. We then followed the same procedure as for local stability to test each metric’s ability to identify a subset of valuable TIMs. For each web we compared the directional determinacy of a model including the selected subset of interaction modifications to a model including just the trophic interactions.

## Results

It was possible to identify the trophic interaction modifications that had particular influence on the system dynamics. In general, those TIMs affecting directional determinacy also affected the local stability of the system. Results for the case of *g*=0.1 and stronger self-regulation terms on producers are shown in Figure 3. Results for other values of body mass-biomass density scaling and different self-regulation terms (SI 3) were broadly similar except in specific cases discussed below. The variability in changes to directional determinacy was related to the median improvement – where a metric tended to give a larger improvement, there was also a greater chance that including those TIMs would result in a reduced model that was considerably worse than not including any TIMs at all (SI 3).

**Figure 3:**
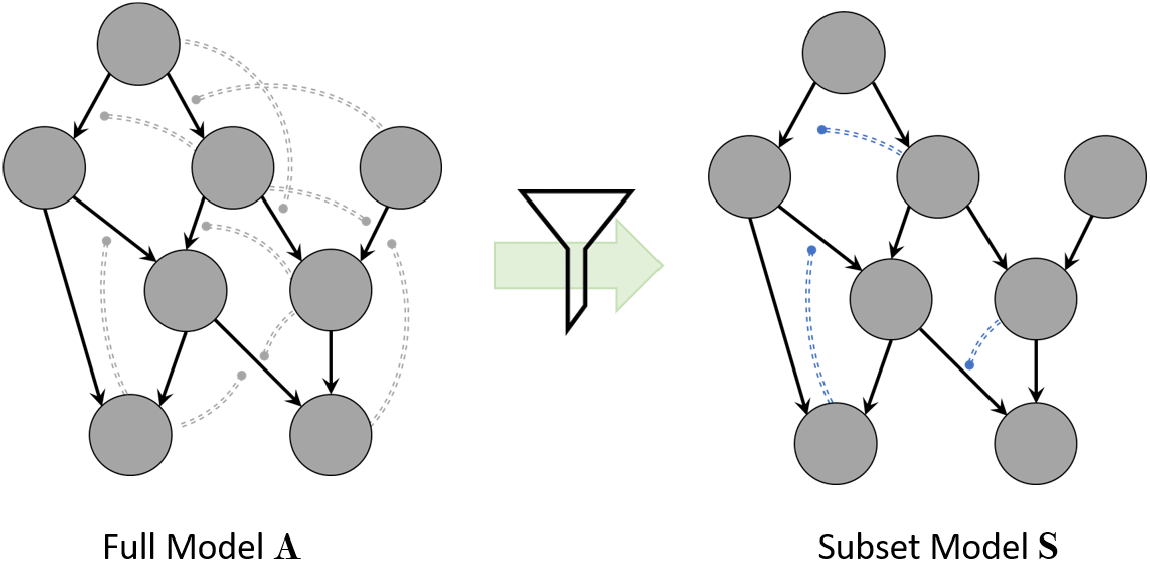
Schematic showing approach to test methods for identifying interaction modifications that are influential in the dynamics of model ecological communities. The trophic interaction modifications in the full community model **A** are quantified and 20% retained using a particular metric to generate a subset model **S**. The capacity for the subset model to represent the dynamics of the full model system are then assessed by their capacity to accurately characterise stability and the direction of response to perturbation as discussed in the main text.

**Figure 4:**
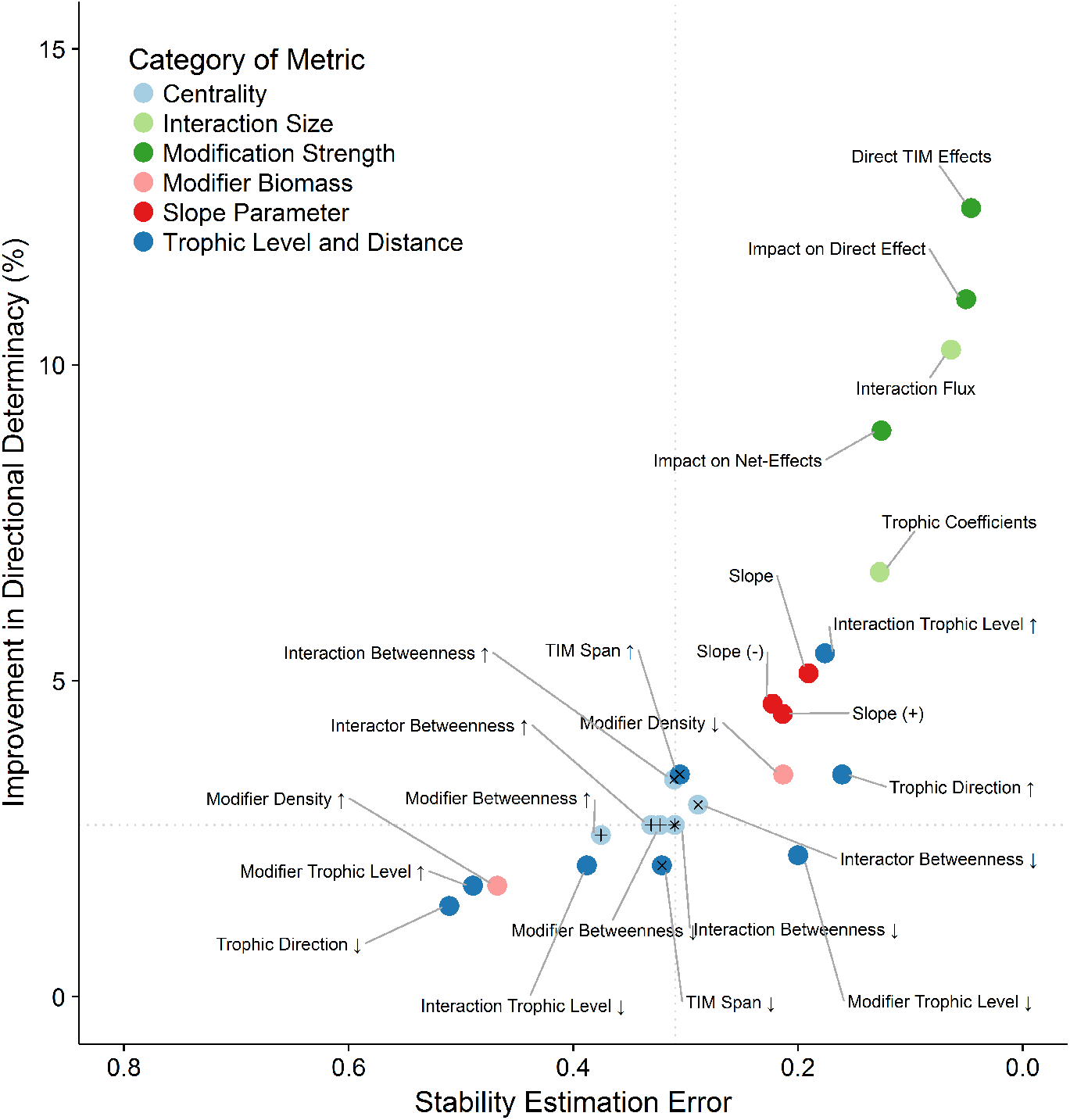
Comparison of the capacity of different metrics to identify TIMs that contribute to the dynamics of whole systems. Arrows indicate whether TIMs were selected for inclusion based on high or low values of the given property. Median error in the estimation of the logarithm of the system stability is plotted on the x-axis, median improvement in the percentage of direction of inter-specific net effects correctly estimated on the y-axes. Grey dotted lines show a baseline when TIMs are chosen at random. Those metrics in the top-right quadrant are therefore better than average at identifying TIMs valuable for both types of dynamics, while those in the lower-left quadrant are worse for both. Points marked with symbols were not significantly different to a random draw for the stability (+), the directional determinacy (x) or both (*). Wilcoxon test (n=2000, *α* = 0.05), paired by underlying network.

In general, those metrics that addressed aspects of the strength of the interaction modification were the most valuable (shown in green in Figure 3) and the very best allowed a near perfect estimation of stability. Metrics based on the centrality of the interaction modifications were the least effective at identifying TIMs, with many not being significantly different to a random draw. Modifications caused by species with a lower biomass density were more valuable than those with high density.

Interaction modifications with large slope parameters of either sign were valuable. However, including just the large strengthening (+ *c_ijk_*) interaction modifications gave a better estimate of the local stability than the strongly negative interaction modifications. This difference was particularly notable for the *g*=0.25 case (SI 3).

The significance of trophic height depended on the relationship between trophic level and biomass density (controlled by the *g* parameter) with modifications of high-trophic level interactions becoming comparatively more important when the webs were more top-heavy. The impact of trophic direction was highly dependent on the distribution of self-regulation terms. Where producers are more strongly self-regulated than consumers, ‘upwards’ TIMs are less influential for directional determinacy.

## Discussion

We have presented methods to quantify the structure of interaction modifications within a trophic network and shown that it is possible to identify properties of trophic interaction modifications that are particularly influential in the dynamics of model communities. This is informative both on a theoretical level in helping to discern how TIMs affect dynamics, but also is useful from a practical perspective in highlighting which TIMs might be most valuable to study empirically. Our work also highlights areas where empirical information is lacking, in particular the need to understand empirical relationships between the strengths of trophic and non-trophic interactions.

TIMs introduce a high level of complexity to ecosystem dynamics. It is therefore reassuring to note that relatively simple and easily observable measures, for example trophic interaction biomass flux, are able to identify the TIMs that drive system dynamics. Similar sets of TIMs were identified as being important in two distinct types of dynamics – response to press perturbations (directional determinacy) and response to pulse perturbations (local stability). By identifying TIMs of high comparative impact we can make progress towards understanding their impact despite the present dearth of data describing the distribution of TIMs in empirical systems. Below we focus our discussion on specific results and their implications, before looking forward to developing this approach in future work.

This study required assumptions regarding the appropriate strength and distribution of TIM values to use. In particular, our results regarding interaction strength are necessarily dependent on the range of the distribution from which the various properties of the model are chosen from. If for instance the range of slope parameters was larger compared to the range of biomass densities, the *c_ijk_* parameter in our model would have been more valuable in comparison to the other TIM strength parameters. Empirical distributions of interaction modification strength in a network do not (to our knowledge) yet exist. Meta-analyses of non-trophic effect strength (Bolnick and Preisser 2005, Preisser *et al.* 2005) suggest an approximate correspondence with trophic interactions. The occasional very strong non-trophic effect at the upper tail of the distribution we used (SI 2) is not necessarily unrealistic as the impact of behavioural effects can in certain circumstances be much faster and stronger than consumptive interactions (Abrams 2001). At the other end of the spectrum, it is necessary to set a threshold where a species’ influence on an interaction is considered too weak to include. This decision trades-off the mean strength and the overall connectance of the modification distribution. An additional complication is how interaction modifications combine to influence an interaction (Golubski and Abrams 2011). For example, interaction modifications may act antagonistically to each other, with combined modifications exerting less effect than would be expected from the individual effects. This would be particularly significant in fully dynamic, non-equilibrium, systems. In our study impacts of this effect are mitigated by our assumption that the trophic interactions are at the post-modification strength.

The increased importance of low-density modifier species via interaction modifications runs contrary to usual expectations for interaction strength distributions. In our study this results from our assumption of the dependence on relative change in modifier populations (Eq. 2). Hence, in our model a change in the density of a low-density modifier will result in a greater modification than the same absolute change in a more abundant modifier. Whether this is a reasonable assumption is to return to the question of how the strength of TIMs is distributed in natural communities. This effect is opposed by trophic interactions being stronger between high-density populations, leading to opposing effects in mixed-process interaction chains. Consider a case where species A consumes species B which then modifies an interaction involving species C. In our model, if B had a lower density, the A-B trophic interaction would be weaker, but this would be more than outweighed by an increase in the B-C non-trophic effect. Further work will be needed to discern how these processes act in nature, but this result highlights that TIMs can greatly increase the importance of species that might otherwise be neglected or dismissed and may disrupt assumptions of the structural patterns of interaction strengths in webs based on trophic interactions only.

Interpretation of the value of topological features of the food web is more challenging as a significant part of their effect can be attributed to correlations with strength-based metrics. In this study we considered topology based on trophic interactions only, as these are currently more widely available and better understood. However, interaction modifications themselves are part of the dynamic topology of TIMs and ideally should be considered in relation to each other.

The most valuable topological metric was the ‘direction’ of the TIM. Those TIMs acting ‘up’ the network were generally more valuable, although the reduced signal with more negative body mass – density relationships suggests this is due to the relationship between trophic height and density rather than pure topological effects. Nonetheless, strongly self-regulated producers may be more resistant to change and hence modifications deriving from them may well be less influential. This highlights that the variability or sensitivity of a species is of equal importance as the magnitude of the modification when considering likely role in the overall system dynamics.

Net-effects between species caused by trophic interactions tend to rapidly decline with the number of trophic links between the interactors (Neutel *et al.* 2002). Where this trophic net-effect is weak, there is a greater potential for a TIM to ‘short-circuit’ the system and dominate the pairwise net-effect between those species. This can be seen in the value of the TIM span metric for directional determinacy, however the comparatively low influence suggests that other significant features of TIMs may be able to override this expectation. Although TIMs have the potential to reduce the dynamic breadth (the number of interactions between the most distant species) of food webs, given that it is estimated that the majority of species are no more than two trophic links apart (Williams *et al.* 2002) the extent of this effect may be limited in practice.

The poor indicative potential of TIM span for local stability, shows that the increased connectance that long-range TIMs introduce has little impact on local stability, as simple models may predict (May 1973). The powerful strengthening TIMs (that benefit the consumer to the detriment of the resource) influence the stability more than weakening TIMs. This is a distinct effect to that caused by mutualism or competitive direct interactions (e.g. Mougi and Kondoh 2012, Coyte *et al.* 2015) since every TIM exerts both a negative and a positive influence. This effect could be driven by at least two mechanisms. Firstly, in our model the assumption of top-down dominated interaction strength imbalance caused by the assimilation term *e* causes non-trophic effects on resources to be larger than on consumers (Eq. 3). This has the result that strengthening TIMs (*c_ijk_*>0) exert larger negative effects (on resources) than positive effects (on consumers). Secondly, there will be a topological effect – strengthening TIMs will lead to consistently positive non-trophic effects on high trophic level species and consistently negative impacts on basal species, which will interact with the underlying tropic interaction distribution.

## Limitations and Future Directions

For this study we made a number of assumptions about the form, topology and strength of TIMs to construct a plausible distribution. The lack of empirical data at a community level is a particular challenge, although identifying non-trophic effects at the network level is possible (Pocock *et al.* 2012, Kéfi *et al.* 2015) and shows that networks of pairwise non-trophic interactions have considerable overall structure (Kéfi *et al.* 2016). Our study emphasises the tight relationship between the underlying trophic network and the consequences of interaction modifications. Isolating an interaction modification from its context, for example in a lab study, will result in the loss of valuable information. The properties of TIMs identified here (Fig. 5) are a first step in building a profile that could be used to identify processes that are more likely to have a considerable influence. Without empirical estimates for the distribution of TIMs it is not yet possible to quantify the potential impact in the manner of Novak *et al.* (2011).

**Figure 5:**
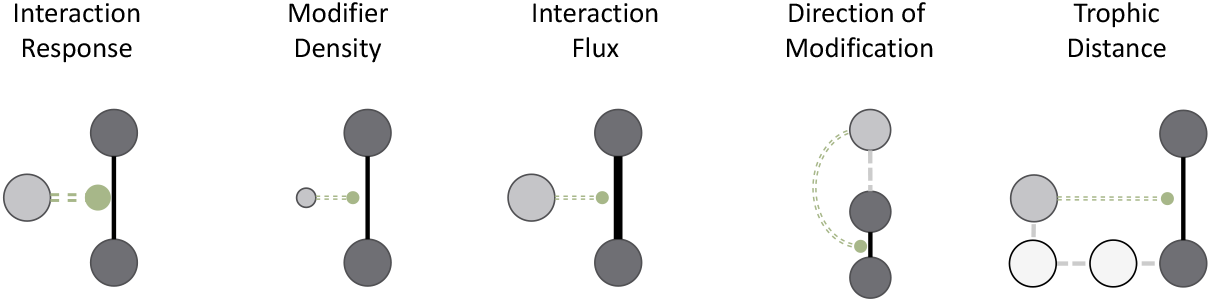
Properties of interaction modifications of relevance to determining their impact.

The instances where directional determinacy actually decreased on the inclusion of TIMs (SI 3) shows the challenges in attempting to improve ecological understanding of complex systems by including additional processes. Our re-sults suggest that there are no easy shortcuts, but that the problem is not intractable. One avenue that will need particular work is the degree of accuracy to which interaction modifications must be known in order to contribute meaningfully to improving ecological forecasts (Petchey *et al.* 2015). Our assumption that all pairwise interaction strengths are known, and that included TIMs are perfectly quantified, constitutes rather wishful thinking for any system of realistic size. Although for small systems, qualitative analyses (Dambacher *et al.* 2003, Dambacher and Ramos-Jiliberto 2007) can be useful for mapping out the potential responses of systems, for larger systems they can rapidly become indeterminate. Exploratory analyses (SI 7) testing the importance in correctly identifying the slope of the TIM function show that as long as the sign of the modification is correct, choosing either random slopes or a single fixed value for the slope can give improvements of mean directional determinacy of 9.8% and 15%, respectively. Future work should more robustly test alternative approximation approaches, such as ‘binning’ effects into broad categories, that have been shown to give good results for estimating trophic interaction strengths (Barabás and Allesina 2015).

Equilibrium studies such as ours allow a precise analysis of particular processes in isolation from other sources of variation. However, this approach does not incorporate the full spectrum of impacts TIMs can have in dynamic systems and future work should seek to address this gap, in particular through fully dynamic models away from equilibrium contexts. Furthermore, trophic interaction strengths are not purely determined by biotic factors – there is also extensive dependence on the abiotic environment (Wootton and Emmerson 2005, Poisot *et al.* 2015). The balance of the impact of exogenous and endogenous sources of interaction strength variation is unknown. The two processes can interact, with TIMs varying depending on abiotic conditions (Sentis *et al.* 2017) and interaction modifications allowing species to resist abiotic variability (Bruno *et al.* 2003). The extent to which interaction strength variation will frustrate ecological prediction is unclear. However, at least some of this variation can be accounted for by dependence on other species in the community which, in principle, is quantifiable (Terry *et al.* 2017).

## Conclusion

To improve our understanding of ecosystems it will be necessary to incorporate important dynamic processes whether they are consumer-resource links, direct non-trophic effects or interaction modifications (Fontaine *et al.* 2011). This potential complexity poses a daunting challenge but not all interaction modifications are equal. We have shown that considering interaction modifications as distinct entities opens up a range of analyses and that although the underlying mechanisms may be complex, relatively simple heuristics may be useful in identifying significant processes in food webs. Where interaction modifications are suspected, but unquantified, our work provides guidance to determine where further research could bring the greatest improvement in understanding. Fully incorporating interaction modifications into the fold of ecology will be a long, but rewarding, endeavour. We hope that our work will encourage further studies, both experimental and theoretical, to understand these critical components of community dynamics.

## Data and Code Accessibility

All code and simulation results are available on the Open Science Framework website: https://osf.io/49jgr/

## Supplementary Material

### 1: Further Details of the Specification of Artificial Communities

#### Defining body masses

Body masses are assigned in a similar manner to Iles and Novak (2016). First, trophic levels calculated on using the pathway method of Levine (1980), rounded into integer trophic levels. These trophic levels were then used to assign body masses were specified on a logarithmic scale (base 10):

i. Producers were assigned a body mass of 0.
ii. Primary consumer (Trophic level 2)) body-masses were drawn from a normal distribution: *N* (0.65, *σ* = 1.52), truncated at 0 to remove the possibility that consumers are smaller than herbivores.
iii. Higher-level consumer (Tropic level 3+) body masses were specified based on the sum of repeated draws from a normal distribution for each trophic level above 2 (h = TL-2), standardising the variance:

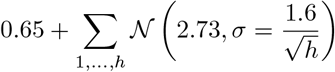

This results in body-masses distributed in a relatively hierarchical manner with trophic level

**Figure 6:**
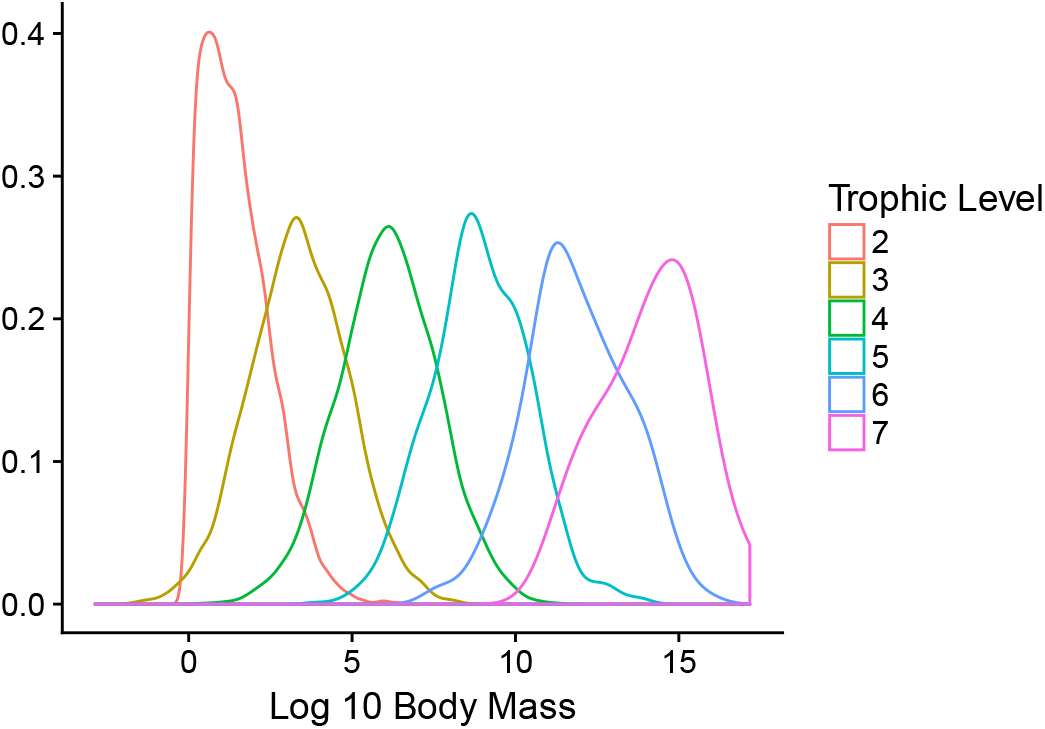
Histogram of body mass distribution with trophic level. Note all Producers have a body mass of 1 (i.e. 0 on the logarithmic scale)

#### Defining attack rates

Biomass density specific attack rates *a_ij_* were set based on documented relationships (Pawar *et al.* 2012) between consumer-resource size ratios and attack rates. We used a scaling exponent (*β*) of 0.95, intermediate between empirical values for 2D and 3D foraging behaviour. We also introduce a generality penalty term term *ω_j_*, which is a simple resource preference term equal to 1/n, the number of resource species of consumer *j*. We don’t include a unit correction term described in the original paper since this would be a constant term in our interaction matrices.

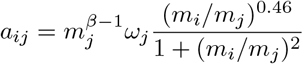

### 2: Distribution of Trophic and Non-Trophic Effect Strengths

**Figure 7:**
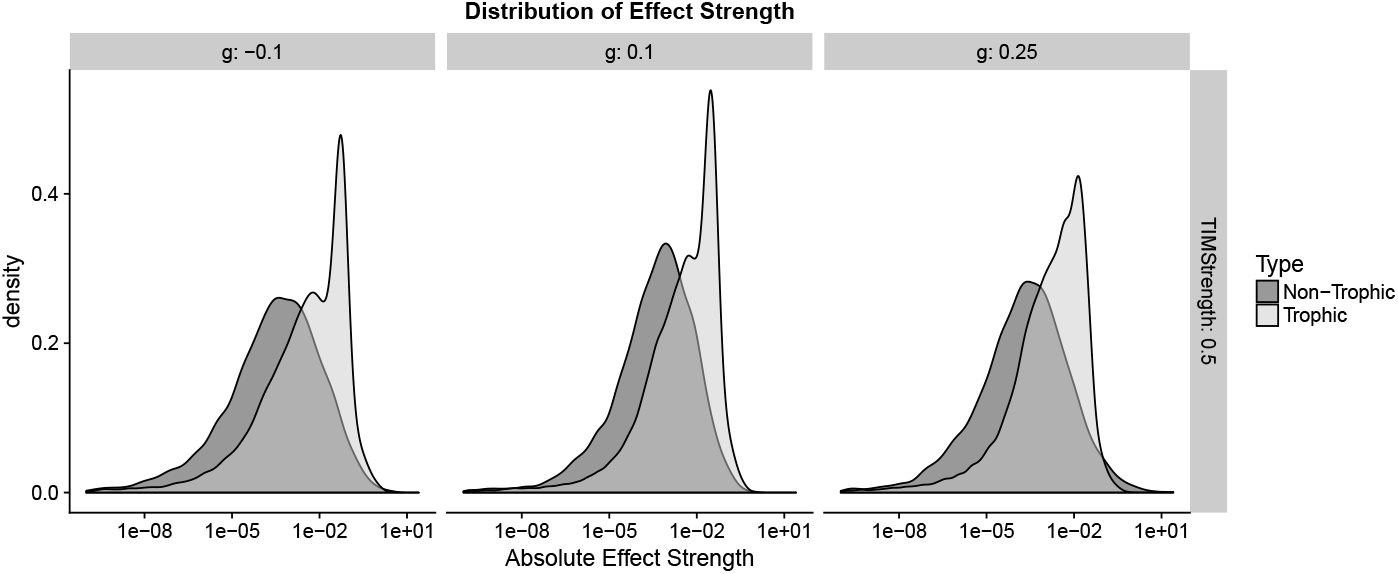
Comparison of the magnitude of the strength of trophic and non-trophic pairwise interaction elements under three different parameterisation of the trophic network.

### 3: Supplementary Results

#### Alternative trophic parameterisations

Figures equivalent to Figure 3 of the main text showing broadly similar results when two properties of the trophic parameterisation are changed - the body-mass:abundance scaling factor, *g*, and the distribution of self-regulation terms, as discussed in the main text.

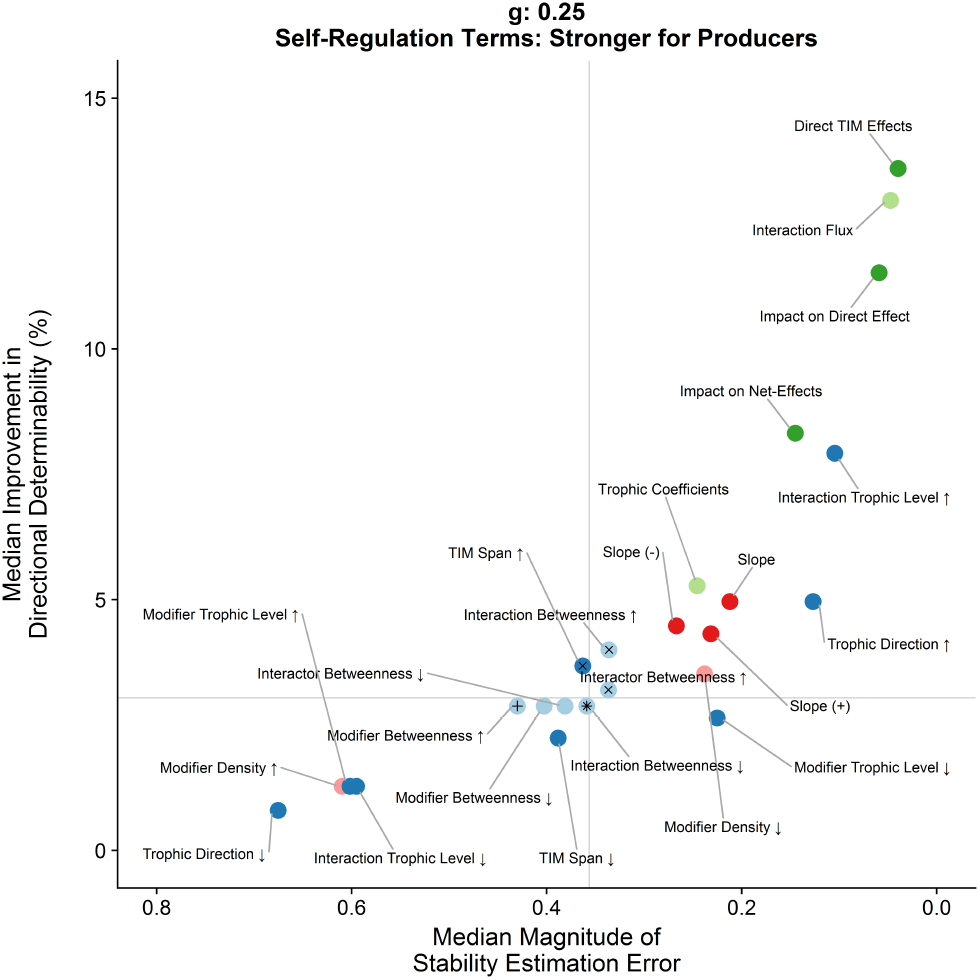

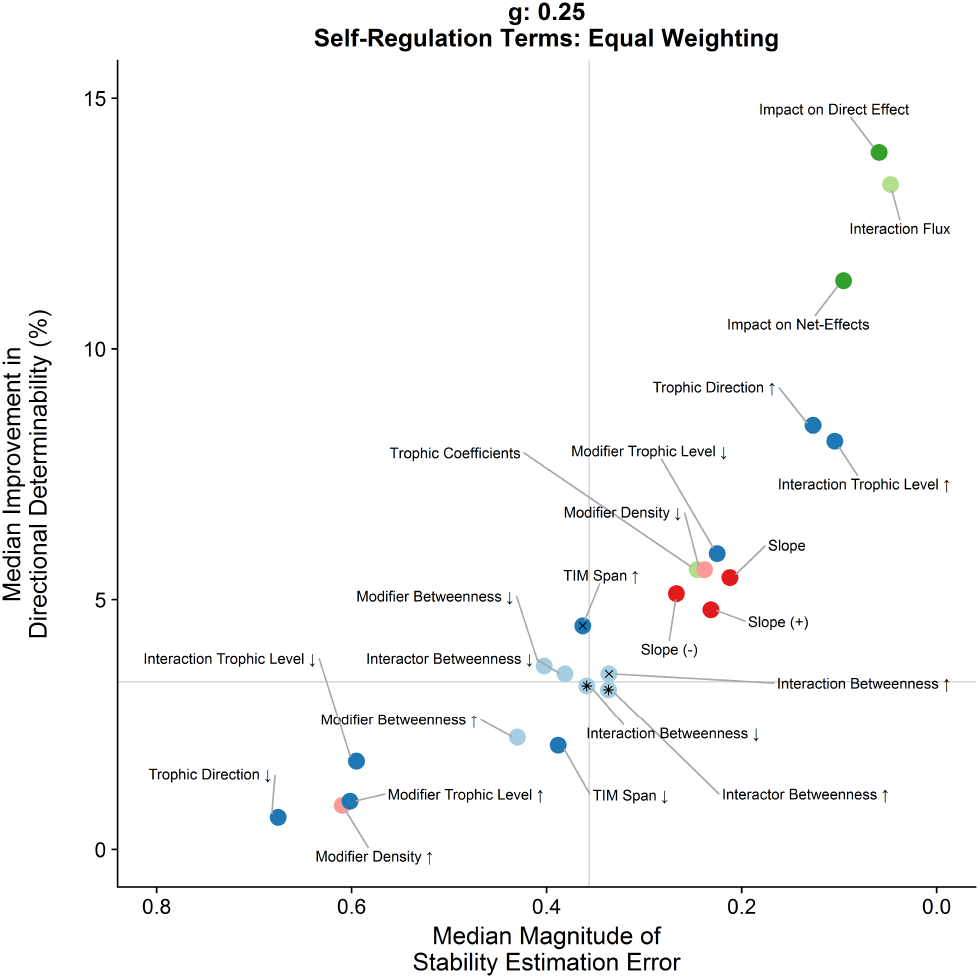

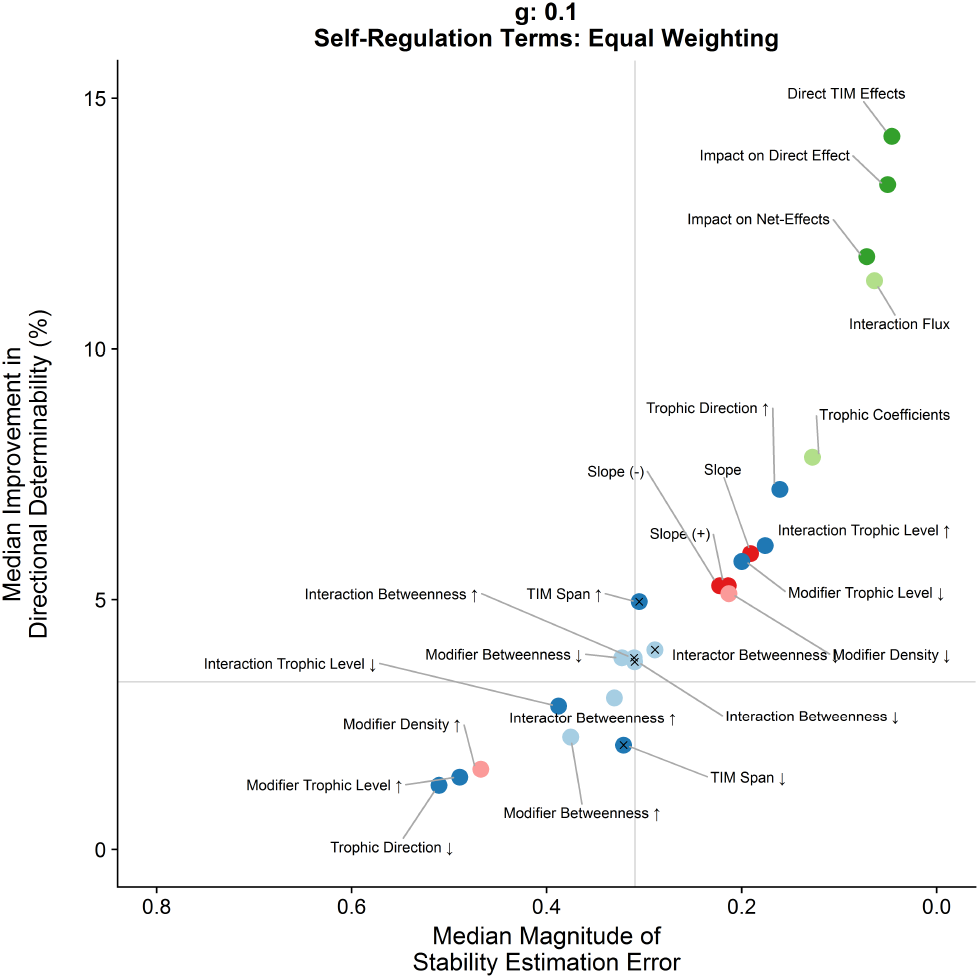

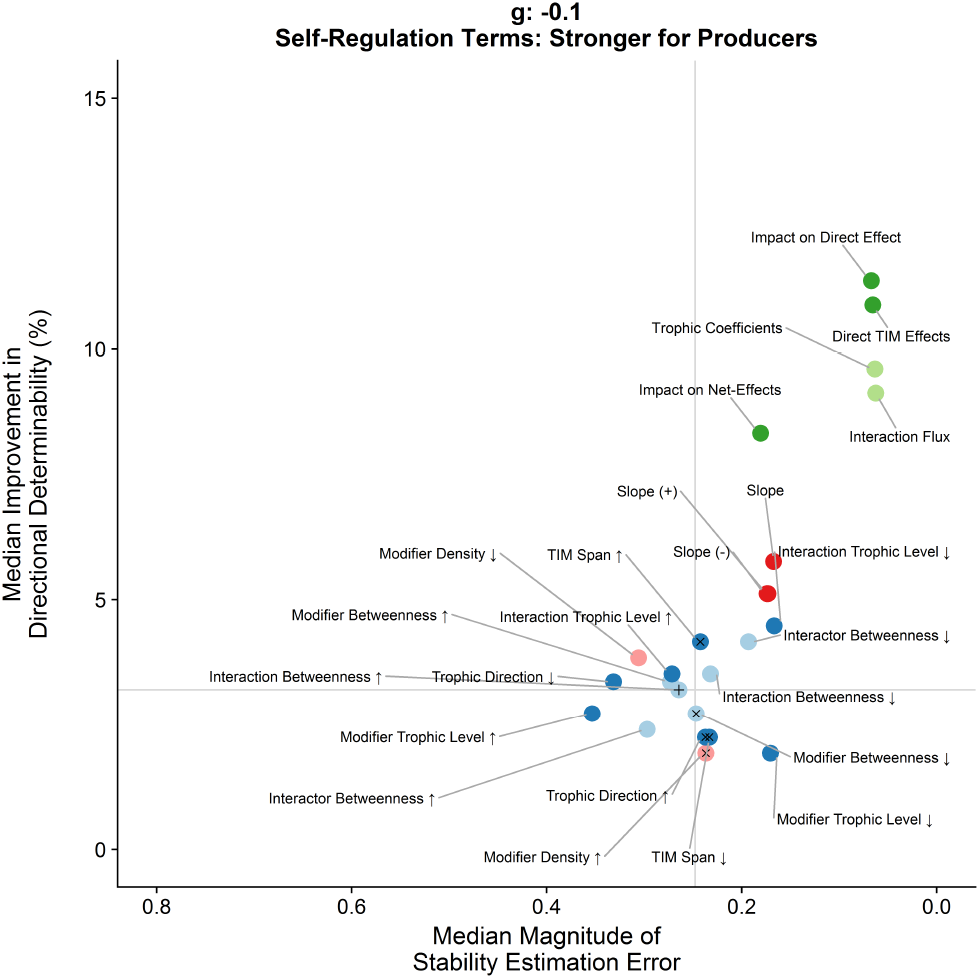

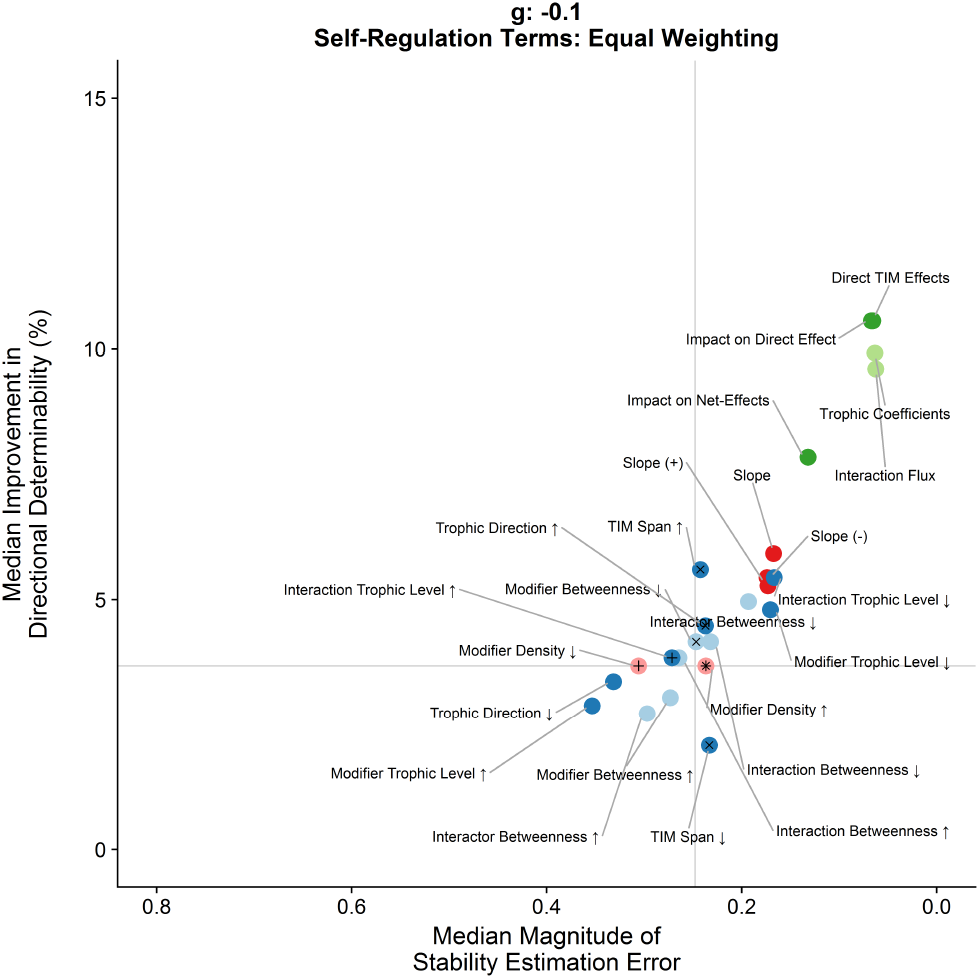

#### Directional determinacy

**Figure 8:**
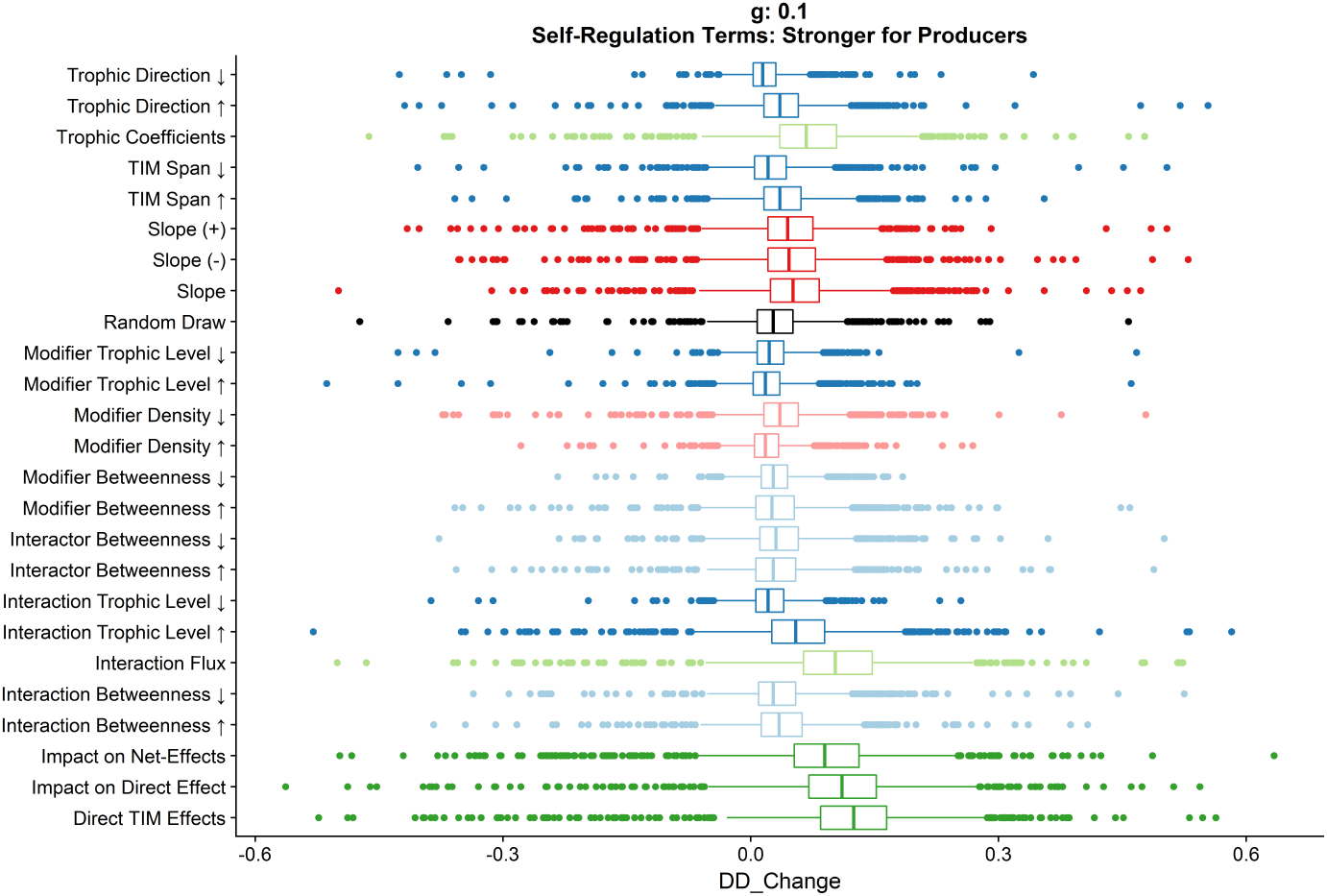
Boxplot showing extent of variance around the median values plotted in the main text figure for each of the metrics in the shift in directional determinacy on inclusion of TIMs as selected by a particular metric

#### Stability estimation

**Figure 9:**
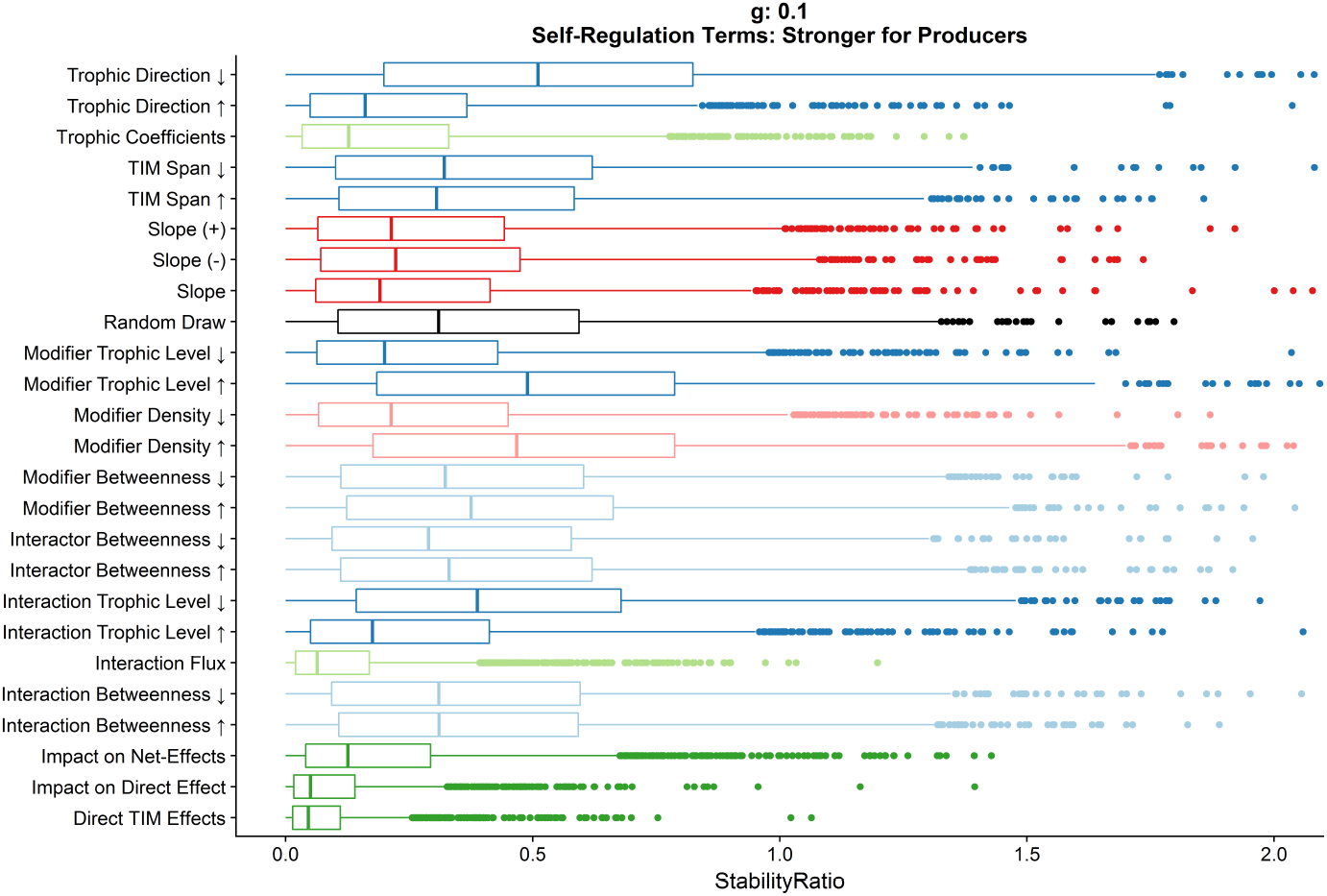
Boxplot showing extent of variance around the median values plotted in the main text figure for each of the metrics in the stability estimation error on inclusion of TIMs as selected by a particular metric

### 4: Condition Numbers of Matrices

The accuracy of numerical inversion of matrices can be sensitive to machine tolerances. The amount of errors that introduced by a matrix manipulation can be described by a *condition number*, often *κ*. A rule of thumb that is often suggested is: If the entries in **A** and **B** are accurate to *r* significant digits, and if the condition number of **A** is approximately *κ*, then the computed solution of **A***x* = **B** should usually be accurate to at least *r −* log_10_(*κ*) significant digits. For ‘normal’ matrices, the condition number for inverting can be directly calculated as the ratio of the largest and smallest eigenvalues 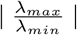. However this does not hold for non-normal matrices, such as our communities, so we calculate *κ* directly using `base::rcond()`.

The figure below shows that although our **A** matrices were relatively ill-conditioned, the numerical accuracy of the inversion was still greater than the vast majority (98.4%) of net-effect elements, in most cases by many orders of magnitude higher. We can therefore be confident of the numerical stability of our results.

**Figure 10:**
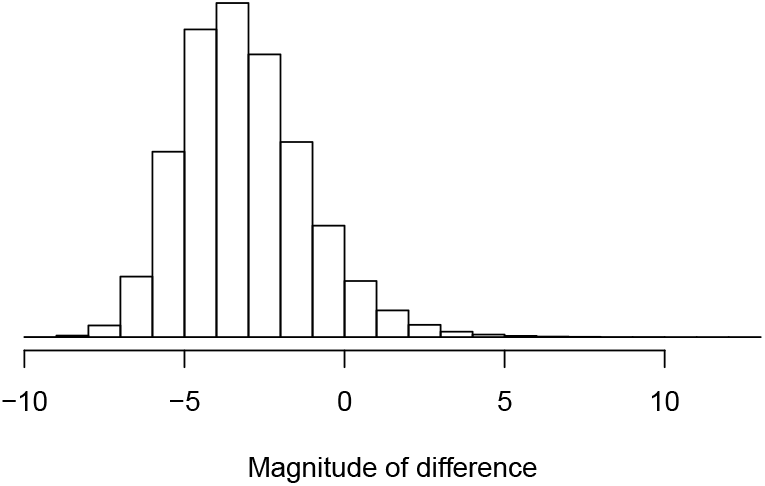
Histogram of differences between reciprocal condition number of interaction matrices and magnitude of net-effect elements

### 5: Statistical Results

**Table 1:**
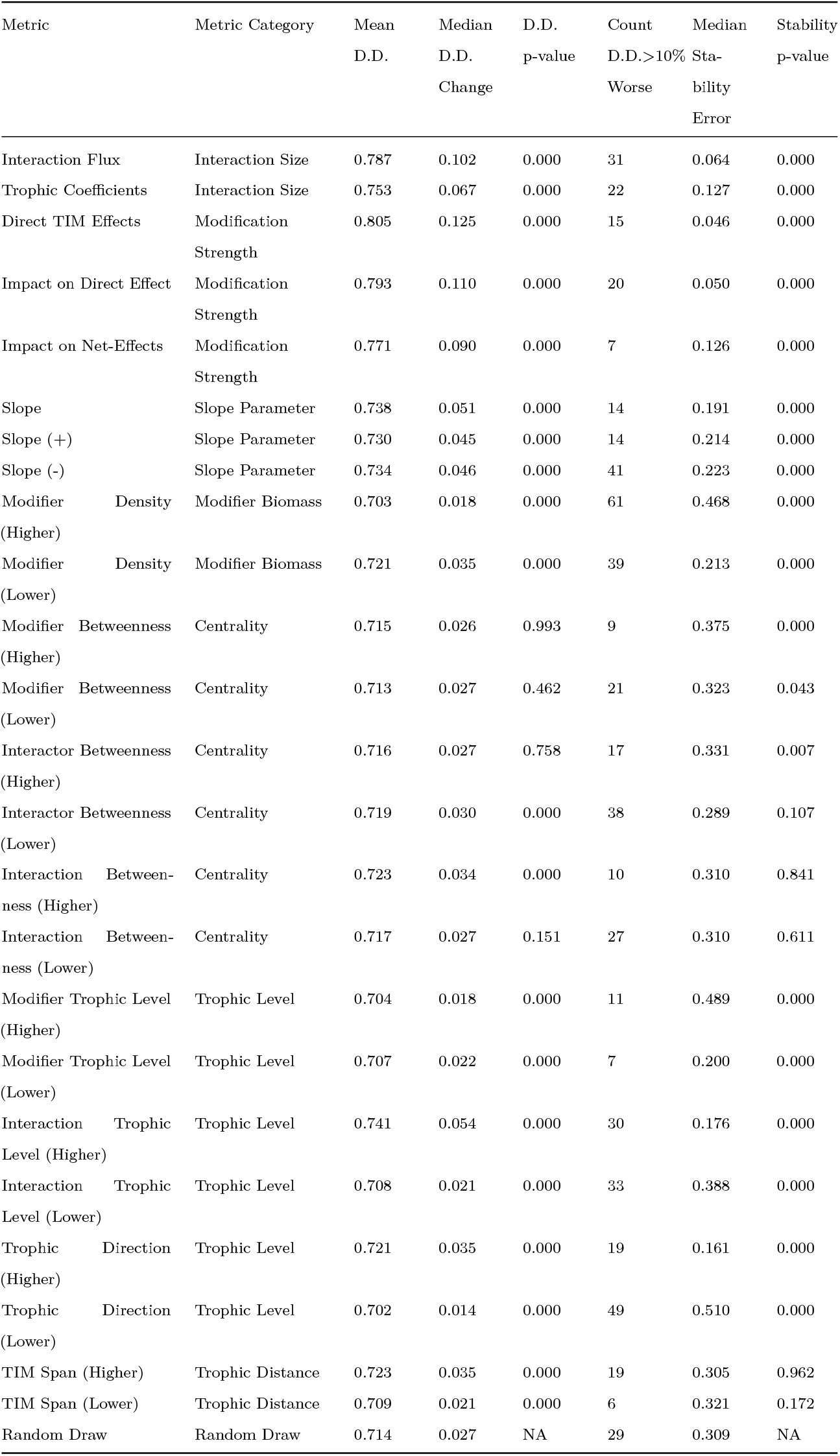
Results tables containing the results of statistical significance tests for other parameterisations are provided as .csv files in the online supplementary material named after the model parameters used to generate them. D.D.= ‘Directional Determinacy’

### 6: Correlation Between TIM Properties for Alternative Trophic Parameterisations

**Figure 11:**
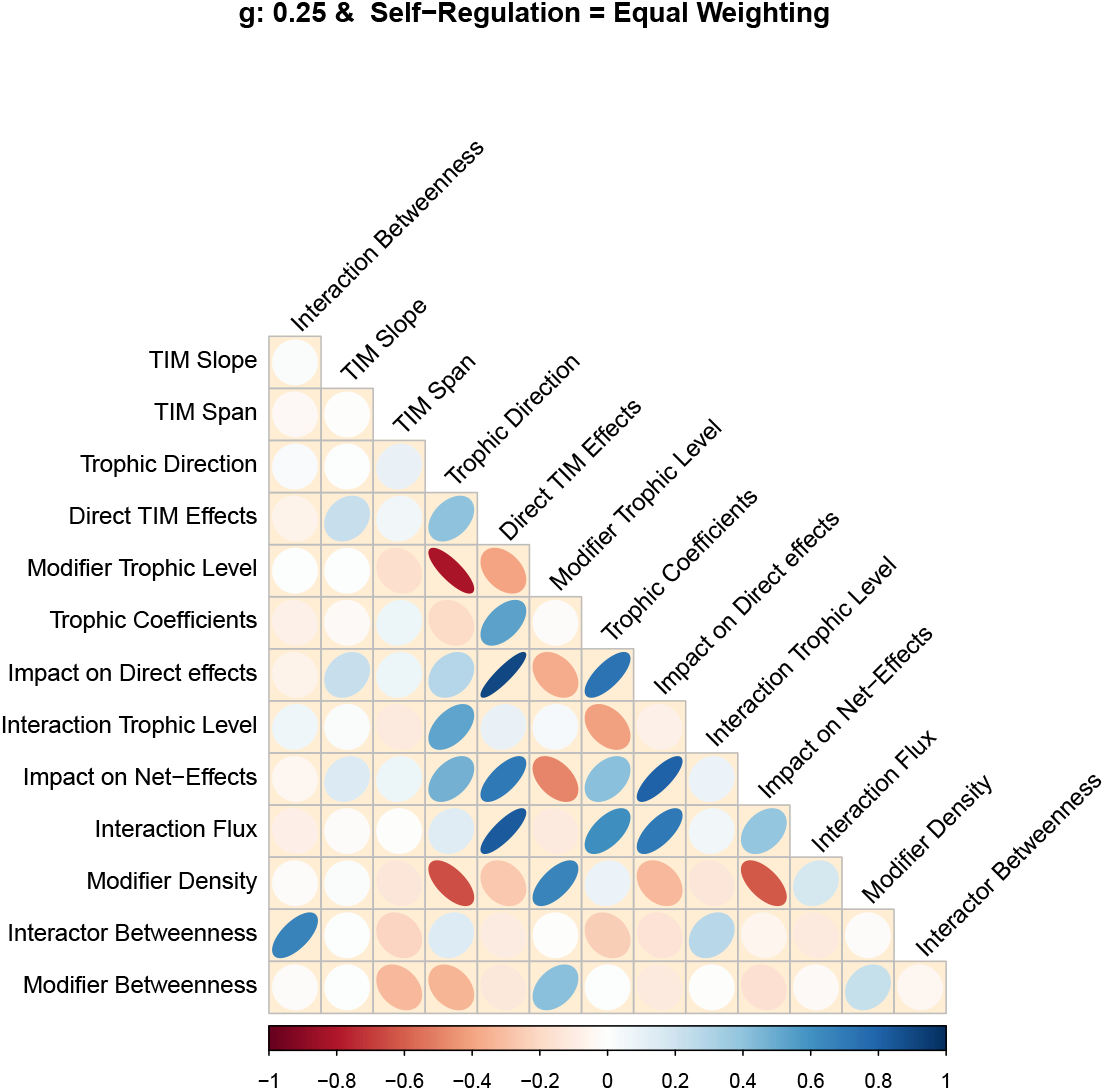
Correlation between TIM properties when the biomass - abundance scaling, g, = 0.25 and self-regulation terms weighted equally for all species

**Figure 12:**
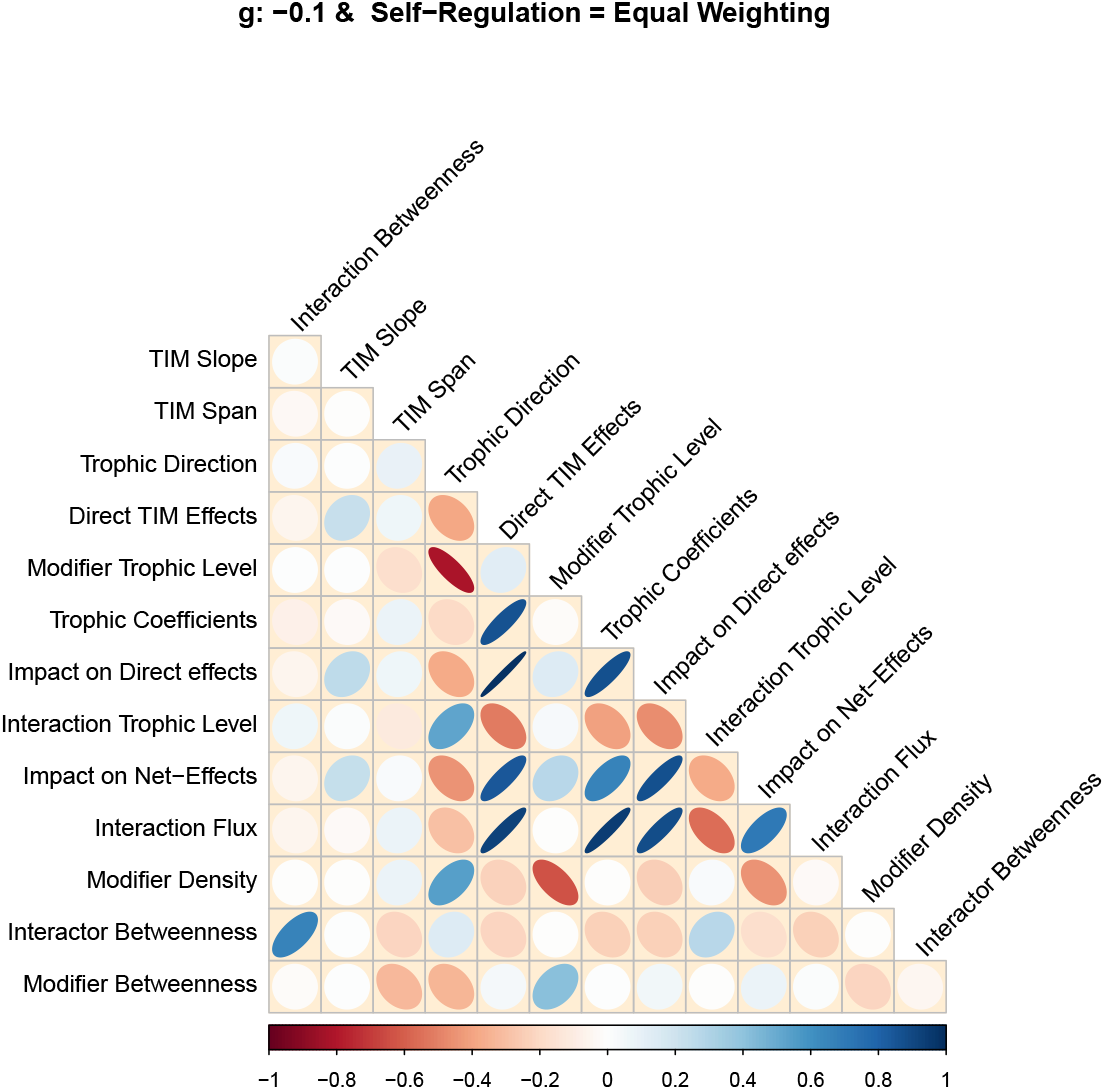
Correlation between TIM properties when the biomass - abundance scaling, g, = -0.1 and self-regulation terms weighted equally for all species

### 7: Exploratory Strength Randomisation Study

To look at the importance of accurately understanding the TIM slope parameters we conducted a simple randomisation study. Using a set of 500 stable communities generated as described in the main text, using *g* = 0.1 and the parameterisation with stronger producer competition.

For each community we conducted two randomisation experiments using the custom function `Randomise_Cijk_Values()`:

1. Re-drawing the *c_ijk_* slope parameters from a uniform distribution, retaining the correct sign
2. Assigning every *c_ijk_* parameter a value of 0.25, retaining the correct sign

In each case the full model and the randomisation model were compared for stability estimation and directional determinacy as in the main text.

| Parameter Values | Directional Determinacy | Stability Estimation |
| --- | --- | --- |
| Random | 0.0978 | 0.1504 |
| Fixed | 0.1553 | 0.1265 |

### 8: Existance and Feasibility of Equilibrium Points

An equilibrium point exists if the net growth rate of each population is zero and can be described as feasible if the density of all of the species at this point is positive (thereby excluding nonsensical cases with negative or zero population densities). In our description of our model communities we focus on the trophic interactions between the populations, directly specify (positive) densities and introduce self-effects according to the trophic position of the species. We make the assumption that the trophic interactions are balanced by additional intrinsic growth or loss terms that we do not directly parameterise since they do not contribute to the off-diagonal terms of the interaction matrices that are the core focus of the paper. Here we demonstrate how these assumptions can be fulfilled in more detail. Historically, analyses of interaction matrix models of ecological communities has been made without reference to the feasibility of the generated communities. Our approach, directly specifying positive equilibrium biomass densities allows us to be assured of the feasibility of our set of model communities.

Interaction matrices of the type we used in this paper (Jacobian matrices) are composed of the elements representing the change to the growth rate of each other species in response to a small change in the density of each species in the community. 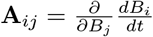. Constant mortality, intrinsic growth rate and migration terms therefore do not appear in the Jacobian matrix. Self-regulation terms only affect the diagonal of the matrix. In order to fulfill the necessary requirement that a feasible equilibrium community has zero-net population change we assume that these constant rate terms are equal to the trophic terms:

Hence:

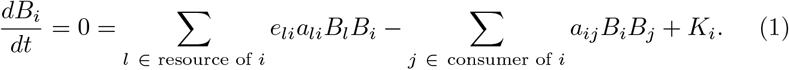

The non-interspecific term *K* can be composed of both constant, linear and non-linear terms: 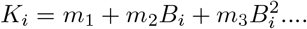 In this formulation the constant terms (*m*_1_) constant inputs such as an external source of immigration, the linear terms *m*_2_ may represent factors such as a constant growth or mortality rate independent of other species in the community, while non-linear terms such as *m*_3_ represent factors such as density-dependent growth or mortality rates. In principle, it would be possible to find algebraically values for *K*, but for simplicity we just discuss them in terms of their signs for each category of species in the community.

For producers, which suffer only negative effects of trophic interactions, *K* would be positive representing intrinsic growth from e.g. photosynthesis. For top-level consumers the *K* terms terms would be negative, representing the dominance of natural mortality terms.

For intermediary consumers the overall sign of the *K* terms necessary to balance the trophic interactions and fulfill our assumption depends on the balance of the trophic interactions affecting each species. In some cases the positive gains from trophic interactions are outweighed by the net losses from consumption by other species in the modeled community. This does not necessarily mean the model is flawed - the positive intrinsic growth required for the equilibrium assumption could derive from a number of processes not included in the basic trophic model used here such as detritivory or immigration.

While the value of *K* does not affect the interspecific interactions, it would have an impact on the diagonal terms of the interaction matrix. The diagonal of the interaction matrix is composed of terms that represent the change in the growth rate of each population in response to a small increase in the population. In studies of interaction matrices there is a long history (e.g. May 1973) of setting these values to constant negative numbers to represent ‘self-regulation’. In biological terms this is often considered to represent increased mortality at higher densities. It is worth noting however that the overall ‘self-regulation’ term is a complex mixture of direct self-regulation terms and terms arising from the trophic interactions of the focal species (see Barabás et al. 2017 for a detailed discussion). For example, in the linear (Holling Type I) functional response model used in this study, an increase in the density of a resource will lead directly to a linear increase in the consumption of the resource and hence a negative effect on the overall rate of population change. Trophic interaction modifications provide an additional mechanisms for ‘self-regulation’ terms. Hence:

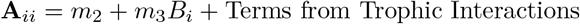

In order to maintain the focus on the interspecific interactions we therefore follow the standard practice of directly specifying the diagonal terms of the interaction matrix. This effectively over-writes the contribution of the *K* term discussed above to the interaction matrix **A**. In our two dynamic response variables, stability and directional determinacy, the diagonal terms only impact the calculation of directional determinacy. In order to test the impact of our assumption of the diagonal terms, we examined two distinct methods of specifying the diagonal terms. Given the lack empirical data on these self-regulatory terms (Barabás *et al.* 2017), we chose these approaches based upon reasonable guesses for the distribution of self-regulation and the requirement that they introduce sufficient self-regulation that over 10% of communities were locally stable.

Approach 1: Each species was assigned a self-regulation term calibrated to the mean strength of the other interactions upon it:

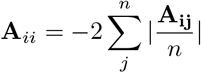

Approach 2: Producers were assumed to be strongly self-regulating while consumers were considered to be weakly self-regulations. Each producer was assigned a term **A**_*ii*_ term equal to twice the strongest trophic interaction in the community while each consumer was assigned a term 100 times smaller.

